# Bioerosion traces and borings in the Upper Devonian vertebrate remains from the Holy Cross Mountains, Poland Bioerosion in fish remains

**DOI:** 10.1101/2021.07.23.453559

**Authors:** Piotr Szrek, Patrycja G. Dworczak, Olga Wilk

## Abstract

Among the hundreds of collected Devonian vertebrate macrofossils in the Holy Cross Mountains, placoderms dominate and provide data on their morphology, distribution and taphonomy. So far 17 out of more than 500 studied specimens have revealed bones with surfaces covered by sediment-filled trace fossils. The traces have been made on the vertebrate remains before their final burial. The borings, oval in cross-section, include dendroidal networks of shallow tunnels or short, straight or curved individual scratches and grooves, which frequently create groups on the both sides of the bones. ?*Karethraichnus* isp. from Kowala and ?*Osteocallis* isp. from Wietrznia are the oldest record of these ichnogenera. Sedimentological clues indicate a shallow water environment, probably from the slope below the storm wave base.

## Introduction

Bioerosion preserved as trace fossils can be biogenic (shells, bones, wood) and abiogenic substrates (rocks, plastic fragments), each representing the results of chemical and/or mechanical processing [e.g. 1–4]. It is essential to understand both the relationships among ancient organisms as well their taphonomy [5]. Bones borings are tools in tracing the different stages of decay as they may record an array of the activity of scavengers [6–7].

This paper reports the bioerosion traces in Late Devonian placoderm bones from the Holy Cross Mountains and elucidate the paleoenvironmental conditions of the Upper Devonian seafloor. Regarding epibionts on placoderms remains, two cases have been reported to date, one from the Middle Devonian of Australia [8] (specimen BMNH P.50313) with the tabulate coral *Aulopora*, and one from the Holy Cross Mountains a supposed bryozoans [9–10]. The latter identification is revised below.

## Geological setting

In the Late Devonian the southern region of the Holy Cross Mountains was located in a subtropical zone at the southern margin of Laurussia with the entire region covered by a shallow, epicontinental sea. An extensive shallow carbonate platform dominated the western part of the region. Its eventual disintegration began at the end of Middle Devonian and resulted in a variety of vast environmental transformations big [11].

Most samples studied come from the Kowala locality (Fig 1), an active quarry located about 15 km south of Kielce at the eastern part of the Gałęzice Syncline. Currently, it represents the most complete Middle-Upper Devonian to lowermost Carboniferous section exposed and examined in Poland [12–22].

**Figure 1.**
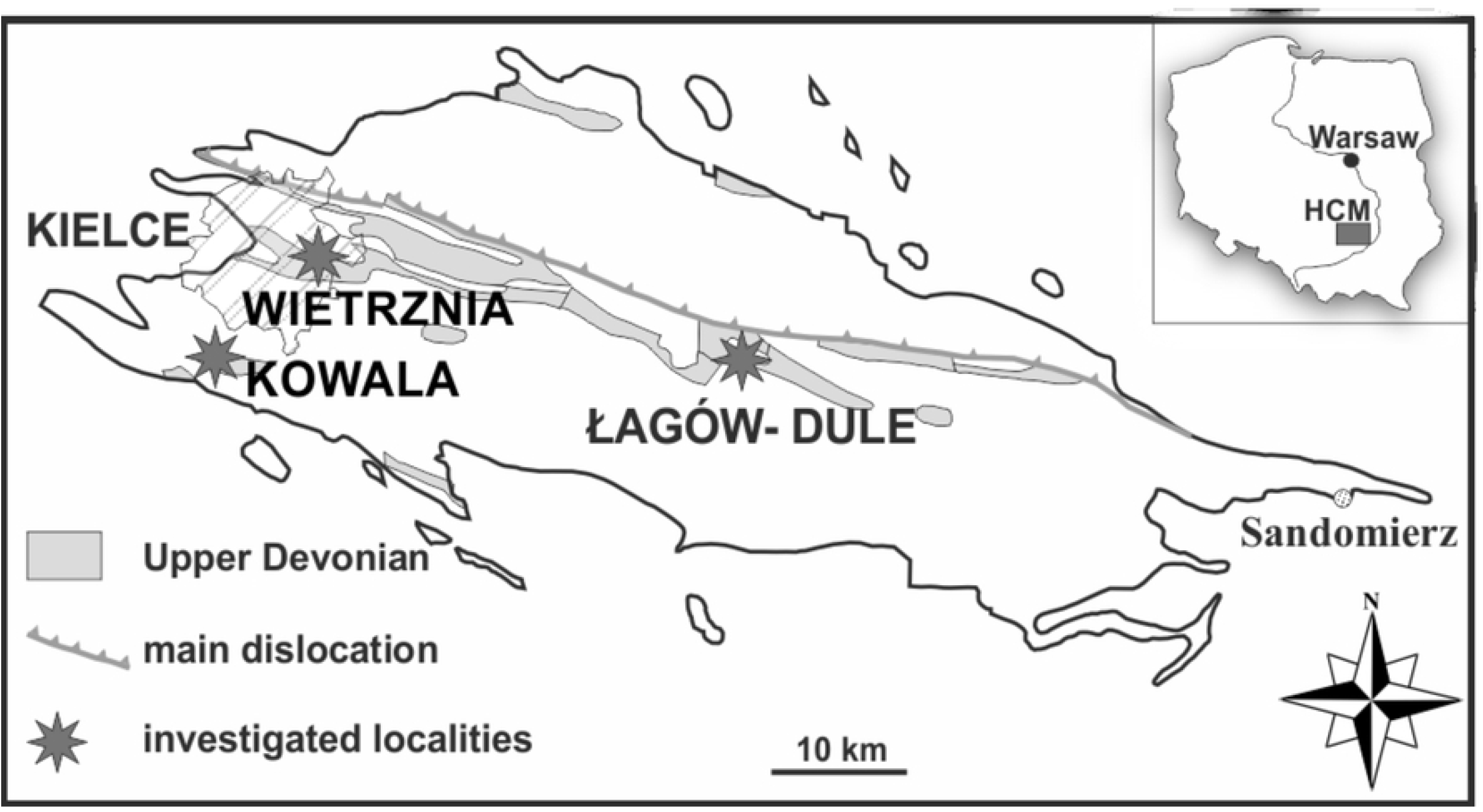
Simplified geological map of Holy Cross Mountains with illustrated Upper Devonian outcrops (adapted from Szrek 2020). The locations of the study areas are indicated by stars.

The second outcrop, the Wietrznia quarry is located in the south-western part of the Kielce region and belongs to eastern part of Kadzielnia Chain, which lies at the southern flank of the Kielce Syncline. The section was a subject of intense research for more than 100 years [23–24]. The Wietrznia quarry has also served as the source of most of the placoderm material studied by Gorizdro-Kulczycka [25] and Kulczycki [26].

All specimens are Upper Devonian in age and were dated based on conodonts. The specimens from Kowala represent the Famennian *Palmatolepis crepida – P. postera* conodont Zone, specimen from Wietrznia is Frasnian, *Palmatolepis falsiovalis* – Early *P. hassi* conodont Zone.

## Materials and methods

Among the hundreds of investigated placoderm bones, 17 have surfaces showing sediment-filled borings. Most of them are poorly preserved, however, 5 bones show features valuable for further research and have been studied in quite some detail. The best preserved traces were found on median dorsal plate (Muz. PGI-NRI 1809.II.29, Fig 2A), an unidentified plate fragment (Muz. PGI-NRI 1809.II.30, Fig 2B), another unidentified plate fragment (MWG UW ZI/43/0070, Fig 2C) and a part of the nuchal plate (Muz. PGI-NRI 1809.II.29, Fig 2D) of an arthrodiran placoderms from Kowala quarry. One specimen is an unidentified dermal bone of a placoderm fish collected in the Wietrznia quarry (Muz. PGI-NRI 1809.II.31, Fig 3). All the bone material has been whitened with ammonium chloride. The specimens are housed in the Geological Museum of the Polish Geological Institute in Warsaw (prefix Muz. PGI-NRI) and at the Faculty of Geology, University of Warsaw (prefix MWG UW).

**Figure 2.**
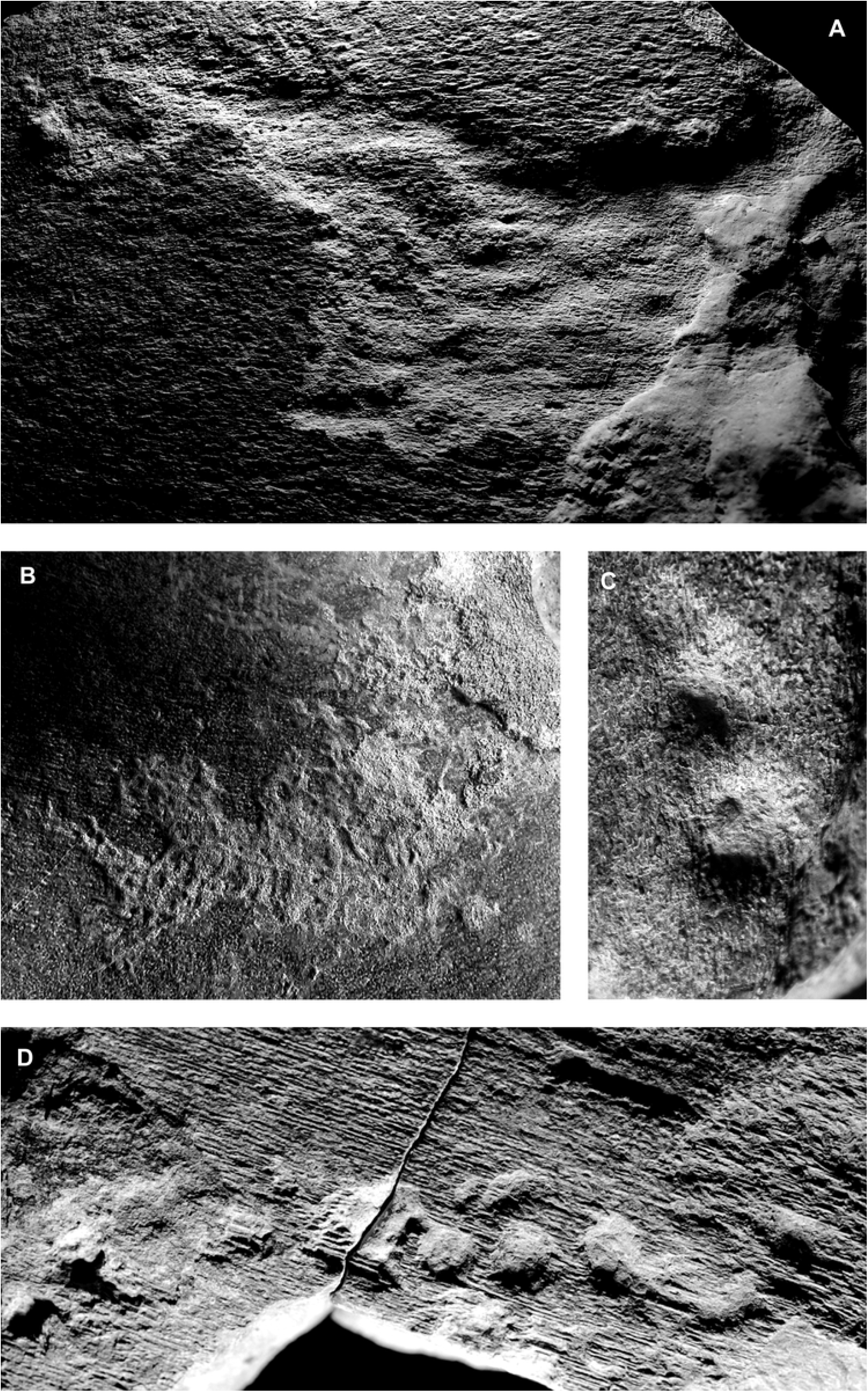
A–median dorsal plate of placodermMuz. PGI-NRI 1809.II.29 showing advanced level of *Arachnostega* sp.; B–unidentified placodermplate fragment Muz. PGI-NRI 1809.II.30 demonstrating ?*Gnathulichnus*; C–external side of unidentified plate fragmentof anarthrodiranplacoderms MWG UW ZI/43/0070 indicating rounded borings *Gastrochaenolites* sp.; D –part of the nuchal plateMuz. PGI-NRI 1809.II.28 showing borings *Gastrochaenolites* sp. visible only on external side of placoderm bone.Scale bar equals 1 cm.

**Figure 3.**
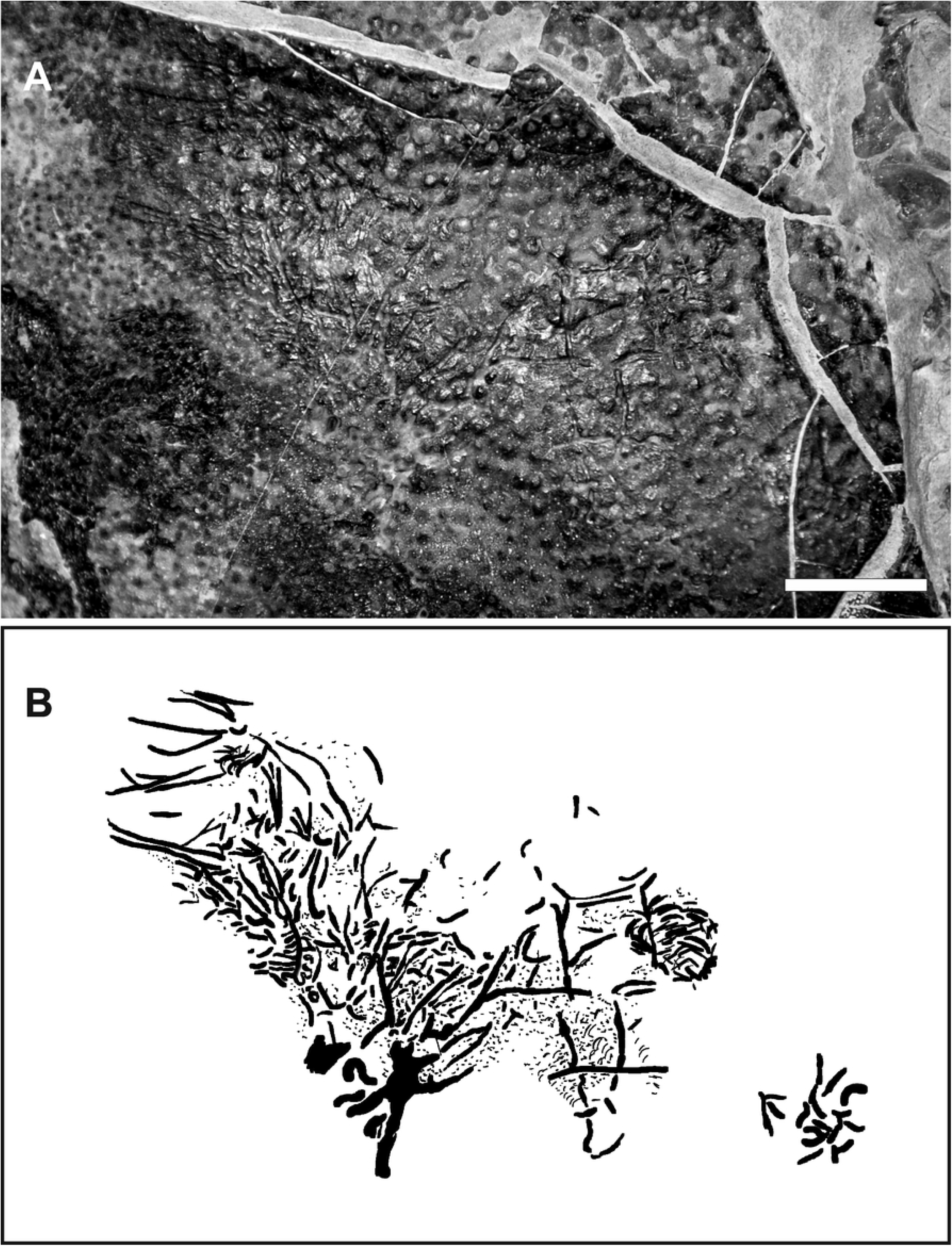
A–dermal bone of a placoderm fish from Wietrznia quarry (Muz. PGI-NRI 1809.II.31) demonstrating long and bifurcate traces and short parallel grazing structuresof herbivorous gastropods and/or polyplacophorans; B–imitative drawing of the traces, ?*Radulichnus* sp. marked by circles.Scale bar equals 1 cm.

## Results

### ?*Osteocallis* isp

Material—Specimen Muz. PGI-NRI 1809.II.30, Wietrznia quarry, Upper Devonian, Frasnian, *Palmatolepis falsiovalis* – Early *P. hassi* conodont Zone.

Description—On the surface of the placoderm dermal bone (Muz. PGI-NRI 1809.II.30, Fig 2B) traces have been preserved as abundant, although irregular grooves. Shallow in depth (0.5-1 mm). Their width varies from 0.4 mm to 1 mm. and the length ranges from 1 to 3 mm. The grooves occur in dense groups and clusters. As they often overlapping other grooves and create more or less star-shaped grooves, the ridges between them are almost invisible.

Remarks—These bioerosional marks closely correspond to *Osteocallis* Roberts et al. [27], which is already known from vertebrate fossils (dinosaur bones from continental deposits of the Upper Cretaceous and the Upper Triassic). *O. mandibulare* shows a network of shallow randomly overlapping trails or paired grooves, which create long path [see Fig 4, 27] and *O. infestans* meandering trail of generally straight overlapping grooves excavated into the bone surfaces. They are randomly orientated and cross each other which are either perpendicular or parallel to one another [see Fig 6, 28].

### ?*Karethraichnus* isp

Material—Specimen MWG UW ZI/43/0070, Kowala quarry, Upper Devonian, Famennian *Palmatolepis crepida – P. postera* conodont Zone.

Description—Rounded borings from 2 to 10 mm in diameter and 3-5 mm depth are visible on the dermal bones from Kowala quarry (MWG UW ZI/43/0070, Fig 2C; Muz. PGI-NRI 1809.II.28, Fig 2D). The precise measurements of the depths of the original borings depth of both specimens is difficult to ascertain due to the amounts of sediment still present on their bottoms as well as from the erosion of the surface of the bone. They terminate in the bone and have rounded, flattened termini. The traces are hemispherical cavities with a circular transversal section and smooth walls. In longitudinal section, they are almost cylindrical-shape and their axis approximately perpendicular to the substrate surface. They appear only on the one side on both specimens and are irregularly arranged on the surface, spaced from each other in distances of 0.5-1 cm increments.

Remarks—The studied borings in the Late Devonian placoderm bones are interpreted as *Karethraichnus* Zonneveld et al. [29]. The first occurrence was described from phosphatic substrate (fossil turtle shells) of the early Eocene. *Karethraichnus* are considerably variable in size and circularity, however *K. kulindros* display deep, non-penetrative wholes. They have similar dimensions: diameter ranges from 0.6 to 5.5 mm and depth ranges from 0.5 to 6 mm. Borings have rounded to flattened, hemispherical terminus. Their longitudinal section is cylindrical and their axis is approximately perpendicular to the surface of bones.

### Morphotypes I, II, III

#### *Morphotype I* (cf. *Radulichnus* isp.)

Material—Specimen Muz. PGI-NRI 1809.II.31, Kowala quarry, Upper Devonian, Famennian *Palmatolepis crepida – P. postera* conodont Zone.

Description—The external side of a placoderm bone (Muz. PGI-NRI 1809.II.31, Fig 3) exhibits not only short, but also slightly curved and shallow structures. They form a cluster of parallel albeit irregular traces. The groups consist of between 5 and 40 individual scratches. Their widths vary from 0.2 mm to 0.4 mm and their lengths range from 0.5 to 3 mm. The scratches often intersected and are cut by another borings.

Remarks—The borings occur on the well-preserved surface of placoderm bone. They are interpreted as *Radulichnus* Voigt [30], grazing traces produced by gastropods or/and polyplacophorans. The ichnogenus has been recognized from invertebrate shells [31], as well as from vertebrate remains [32, Plate 2, Fig 1–2]. The borings exhibit both straight or curved scratches which appear to be parallel, subparallel or irregular and form different morphological patterns (meandering line, patches or grooves). Until now, two ichnospecies are known: *Radulichnus inopinatus* Voigt [30], consisting of tight bundles of 4-6 subparallel, rather straight scratches, and *R. transverses* [31], which more often forms lines rather than rows, and frequently a wider spacing between individual furrows. The studied arched and short traces create groups of cluster more abundant than *Radulichnus inopinatus* and they not cover all surface of the substrate.

#### *Morphotype II* (cf. *Palaeomycelites* isp.)

Material—Specimen Muz. PGI-NRI 1809.II.31, Kowala quarry, Upper Devonian, Famennian *Palmatolepis crepida – P. postera* conodont Zone.

Description—This specimen was interpreted by Szrek [9–10] as a supposed bryozoan etching traces, but its morphology, especially regarding no multiple ramifications and smaller diameter of single burrow (Fig 3). The shallow, irregularly arranged cylindrical galleries are usually short and irregularly contorted, rarely groves are long and bifurcate. Their width varies from 0,2 mm to 0,4 mm and their length ranges from 3 to 20 mm. They occur on a sizable area of the plate with one group consisting 20 individual borings 3 mm long. Slightly curved, the traces follow a more or less straight course with respect to the plane of the substrate, displaying strong curvature as well as sporadic branching.

#### Morphotype III

Material—Specimen Muz. PGI-NRI 1809.II.29, Kowala quarry, Upper Devonian, Famennian *Palmatolepis crepida – P. postera* conodont Zone

Description—Specimen no. Muz. PGI-NRI 1809.II.29 (Fig 2A) is covered by dendroidal networks (meshwork) of shallow tunnels, oval in cross-section. They are visible as smooth furrows forming significantly and dense networks. The network continues laterally, ramifying and branching consequently. The width of an individual varies from 0.4 cm to 3.5 cm and their depths ranges from 0.5 mm to 1 mm. There is no visible reduction of diameter at the ramifications, and the ridges between them are almost invisible. In the central parts of structures, the traces compose areas of overworked surface with details of single burrows poorly visible. These patterns of the net-like burrow scan possibly indicate that the specimen Muz. PGI-NRI 1809.II.29 demonstrated the advanced stages of the growth and possible escalation of bioerosion.

## Discussion and conclusions

Bones of placoderms with epibionts or borings have been reported only by Bystrow [33], White [8] and [Szrek 9–10]. Judging by the numbers of bones studied here, one may summarize that this low number of finds has resulted from the lower level of the attention has been paid to this phenomenon. Contrary to previously suggestions, traces of bioerosion may be present on a considerable number of specimens. During the Silurian and Devonian diversity of bioerosion has increased but has hardly been recorded from vertebrate remains.

Our material provides clear examples of bioerosion in the Late Devonian vertebrate remains from the Holy Cross Mountains. Beyond doubt, all studied specimens have been bored *post mortem* either by scavengers or grazing organisms because all have been attacked from both surface sides. Hitherto, *Karethraichnus* was recognized as *syn vivo* interaction [29]. Affected turtles bones display evidence of healing of the borings, what indicate parasitic relationship. In the presented specimens we did not observe any malformation of bones, which could demonstrate *syn vivo* interaction between placoderms and the trace marker.

The specimen from the Kowala quarry has been interpreted by Szrek [9–10] as bryozoan etching traces provisionally attributed to hederellids (?). Bryozoan colony of should develop only on surface above the sediment unless the colony settled one in case of reverse of the specimen, the second side. The environment however, does not support any rotation due to the low-energy conditions identified in this part of the section.

The studied placoderms bones display the oldest evidence of ?*Osteocallis*. The bioerosion differs from *Gnathichnus* Bromley [34] and *Radulichnus* Voigt [30] by having arcuate, paired scratches, which can create paths of scratches or cross each other. We also demonstrate the oldest example of ?*Karethraichnus*. These deep cylindrical borings differ from ichnogenus *Gastrochaenolites* (carbonate substrate) and *Teredolites* (wood) of the substrate in which they occur [35].

On the placoderm bones grazing structures of herbivorous organisms have been recognized. The bones represented the only hard substrate for microbial mats or algae. The distribution of these grazers depends on an occurrence their food. Hence *Radulichnus* characterizes shallow water environments [34, 36–40]. But deeper environments cannot be exclude, where fish remains could be covered by microbial mats. Since organisms which left *Radulichnus* feed on algae and microbial mats, the patter of furrows depend on their distribution on the substrate. Thus the isolated patches reflect the surface of placoderm bone covered by algae. Furthermore, the groups are similar to borings made by *Gibbula* sp. illustrated by Thomson et al. [36] and Gibert et al. [41], which create groups of shallow, short and curve scratches. Additionally, we observe 2 sizes of these trace, indicating that organisms grazed at different time on the placoderm bone.

The environment had to stable enough to provide object a steady position on the seabed as the sedimentation ration must have been low [42]. Large plates of placoderms could provide an area of development of microbial mats, which the generally soft, muddy sea-floor as a secondary hard substrate. However, big sizes of vertebrate remains did not cause that they were the most common encrusted organic substrates during the Devonian. The traces from the Holy Cross Mountains most likely resulted from opportunistic behaviour of the trace maker boring, as the bones could have represented the only hard elements at the sea floor.

## Acknowledgments

We are grateful to Michał Poros (Geopark Kielce) for his help in providing part of the specimens presented herein. PS and OW were financed by the Polish Geological Institute-National Research Institute (grant no. 62.9012.1916.00).

